# The impact of isoniazid preventive therapy on tuberculosis among household contacts of isoniazid–resistant patients

**DOI:** 10.1101/479865

**Authors:** Chuan–Chin Huang, Mercedes C. Becerra, Roger Calderon, Carmen Contreras, Jerome Galea, Louis Grandjean, Leonid Lecca, Rosa Yataco, Zibiao Zhang, Megan Murray

## Abstract

**Background:** The World Health Organization recommends the use of isoniazid (INH) alone or combination INH and rifapentine therapy to treat latent tuberculosis infection (LTBI) in groups at high risk of tuberculosis (TB) progression. The recent rise of INH– and multi–drug resistant (MDR) TB has complicated the choice of LTBI treatment regimen. We examine the risk of TB disease among household contacts (HHCs) who received INH after being exposed to patients with drug–sensitive, INH–resistant, or MDR tuberculosis.

**Methods:** In this prospective cohort study conducted Between September 2009 and August 2012 in Lima, Peru, we identified 4,500 index TB patients and measured incident TB disease in their 14,044 HHCs over a one–year follow–up period. HHCs under 19 years of age were offered INH preventive therapy (IPT). We used a Cox frailty proportional hazards model to evaluate whether the effect of IPT on incident TB disease varied by the resistance profile of the index case. We repeated the analyses in a second independent dataset.

**Findings:** Among 4,216 HHCs under 19 years of age, 2,106 (50%) initiated IPT at enrolment. The protective effect of INH was more extreme in HHCs exposed to drug–sensitive (Hazard Ratio [95% confidence interval]=0·2[0·20–0·50]) and to MDR–TB (0·26[0·08–0·77]) compared to those exposed to mono–INH–resistant (0·80[0·23 to 2·79]). Among those who received at least three months of INH, effectiveness increased across all three groups (INH–sensitive:0·20 [0·10 to 040]; MDR:0·16 [0 02–127]; mono–INH–resistant:0·72 [0·16–3·16]). In the second independent study, TB occurred in none of the 76 HHCs who received IPT compared to 3% (8/273) of those who did not.

**Interpretation:** We found that IPT use is associated with reduced incidence of TB disease among contacts of MDR–TB patients. This finding suggests that INH may have role in the management of MDR–LTBI.

**Funding:** National Institutes of Health and the National Institute of Allergy and Infectious Diseases CETR (U19AI109755) and TBRU (U19AI111224)

**Research in Context:** *Evidence before this study:* Few data exist on the efficacy of INH in preventing TB progression among people exposed to MDR-TB. In a study from Brazil, researchers reported that the risk of TB disease among 190 TST–positive contacts of MDR–TB patients was 2·3 times lower among those who received IPT than among those who did not. In Israel, investigators reported no cases among 71 contacts of MDR patients who received IPT during a six–year follow–up period. In South Africa, researchers reported that children who did not receive any preventive therapy were four times more likely to develop TB disease than those who received less than six months of an individualized preventive therapy regimen that contained high dose INH (15–20 mg/kg/d). Several other studies that reported on INH in contacts of MDR-TB patients lacked control arms and thus the efficacy of INH could not be measured.

*Added value of this study:* Here, we found that IPT protected contacts of INH–resistant TB patients from developing TB disease. Isoniazid preventive therapy effectiveness was increased among contacts who received more than three months of treatment and under five–year–old.

*Implications of all the available evidence:* Our findings suggest that INH may have a role in the management of MDR–LTBI

## Introduction

The worldwide TB pandemic remains one of today’s greatest global health challenges. The World Health Organization (WHO) estimates that there were 10·4 million new cases of TB in 2016 and that one quarter to one third of the world’s population has latent TB infection (LTBI).^1,2^ Although treatment of LTBI has been shown to protect against TB disease progression, only a tiny minority of those at risk receive preventive therapy.^2^ WHO’s recently revised guidelines on treating LTBI now recommend systematic testing and treatment of LTBI for an expanded group of people at high risk of TB disease including child and adults contacts of pulmonary TB patients. Recommended regimens for LTBI include six to nine months of isoniazid (INH), a three–month regimen of rifapentine plus INH, three to four months of INH and rifampicin, and three to four months of rifampicin alone.^2^

The recent rise of INH–resistant and multi–drug resistant TB has complicated the choice of an LTBI treatment regimen. Although several small studies have shown that regimens tailored to specific drug–susceptibility profiles can be effective, most of these lacked control arms or else compared these individually tailored regimens to no treatment rather than an alternative regimen.^3^ WHO concludes that the current lack of evidence on optimal regimens prevents the formulation of definitive recommendations for INH–resistant and MDR–exposed contacts.^2^

In countries that implement preventive therapy for those at high risk, close contacts of MDR–TB patients often receive standard LTBI regimens prior to time that the index patient’s drug susceptibility tests are available to the treating clinician. In areas where rapid diagnostic tests for MDR are not yet available, contacts may receive INH for months prior to the eventual diagnosis of MDR.^4,5^ Here, we examined the risk of disease progression of individuals exposed to sensitive, INH or MDR–TB who received Isoniazid preventive therapy (IPT) as part of routine TB management.

## Methods

### Recruitment

This study was conducted in Lima in 106 district health centers that provide care to a population of approximately three million residents. Patients were referred to study staff if they were over 15 years of age and had been diagnosed with pulmonary TB (PTB) disease by a health center clinician. We requested permission to visit each patient’s household and recruit his or her household contacts (HHCs) into a prospective cohort study. Study workers aimed to enroll all household members within one week of the diagnosis of the index case.

### Baseline assessment of index patients and household contacts

We collected the following data from index patients and HHCs at the time of enrollment: age, height, weight, gender, occupation, history of TB disease, alcohol, education, housing information, intravenous drug and tobacco history, symptoms of TB, BCG vaccination, and comorbidities including HIV and diabetes mellitus. For index cases, we collected the duration of symptoms before diagnosis, presence of cavitary disease, sputum smear status, and culture results. Those with positive cultures had drug–susceptibility tests and MIRU–based genotyping (detailed information in Supplement 1). For HHCs, we noted whether IPT had been initiated. HHCs with symptoms of TB disease were referred to their local health clinic for chest radiography and clinical evaluation. HHCs with no known history of TB disease or previously documented infection received a tuberculin skin test (TST).

### *INH preventive therapy* for HHCs

The 2006 Peruvian National TB Program recommended that HHCs 19 years old or younger and those who had a specified comorbidity should receive six months of IPT while those with HIV should receive 12 months.^6^ Children aged 19 and under were offered IPT at the time index patients were diagnosed, regardless of TST status. Health care providers often chose to discontinue IPT in HHCs if the index patient was subsequently diagnosed with MDR–TB but some MDR–exposed HHCs received a full course of IPT. We used medical records from participating hospitals and health clinics to determine the duration of IPT.

### Follow-up of household contacts

Participants were reassessed at two, six, and twelve months; those with TB symptoms were referred to their local health center for further clinical evaluation including a chest radiograph and sputum smear.

### Outcome definition

We identified incident TB among HHCs during scheduled household visits and from a systematic review of TB registries at the participating health clinics. We considered HHCs to have co–prevalent TB if they were diagnosed within two weeks of the diagnosis of the index patient and to have secondary TB otherwise. Diagnosis of TB among adult HHCs followed the same criteria as outlined above. We defined TB disease among contacts younger than 18 years of age according to the consensus guidelines for classifying TB disease in children.^7^

### Analyses

We restricted the analysis to HHCs under 19 because older contacts were offered IPT only if they had comorbidities that substantially increased their risk of TB disease. We used a Cox frailty proportional hazards model to evaluate risk factors for incident TB disease, accounting for clustering within households.^8^ We first performed a univariate analysis to examine the effect of IPT on TB incidence, followed by a multivariate model adjusting for index case age and the age, social economic status (SES), and TB history of the HHC. To evaluate whether the effect of IPT on TB incidence varied by the index patient’s resistance profile, we added a variable representing index–patient INH–resistance and an interaction term for INH–resistance and IPT. Because the spectrum of INH–resistance causing mutations that lead to INH mono–resistance may differ from those that lead to MDR–TB, we classified strains as sensitive, mono–INH–resistant, or MDR–TB (resistant to both INH and RIF). Previous studies have shown that the effectiveness of IPT treatment is reduced if the treatment duration is less than three months.^9^ We therefore repeated these analyses stratifying on duration of treatment. We also considered the possibility that HHCs ≤ five years of age would be more likely than older participants to acquire TB at home rather than in the community and so conducted sensitivity analyses restricted to this subgroup. To determine whether the effect of IPT was related to the mean inhibitory concentration (MIC) of the infecting organism, we repeated these analyses for the subset of HHCs exposed to index patients for whom quantitative INH–resistance was available.

### Verifying our finding with an independent dataset

We conducted a separate analysis using publically available data from a prospective cohort study in South Lima and Callao, Peru between 2010 and 2013, posted by Grandjean et al.^10^ This study enrolled 1,055 HHCs of 213 MDR–TB index cases and 2,362 HHCs of 487 drug–susceptible index cases and measured incident TB over two–years of follow–up. Drug susceptibility testing for INH and RIF was performed for all index cases’ samples using microscopic observation drug susceptibility assays in regional laboratories and results were confirmed in the national reference laboratory using proportions methods.^11^ The investigators note that IPT was discontinued in this group after MDR–TB index cases were confirmed but data on the duration of IPT were not available. We applied the same analytic plan which we used for our own data to this independent dataset.

## Results

### Data collection

We enrolled 14,044 HHCs of 4,500 patients suspected of having TB, of whom 12,767 had been exposed to index patients with microbiologically confirmed TB. Of these, 5,496 (43%) were 19 years of age or under. We restricted our analyses to 4,216 HHCs who were exposed to an index case whose INH resistance profile was available (Figure 1). At the time of enrollment, 2,106 HHCs (50%) had initiated IPT. On average, the duration of IPT was shorter among HHCs of MDR–TB cases (115 days) than those of drug–sensitive TB (142 days) and mono–INH–resistant TB cases (148 days) (Figure 2). The baseline characteristics stratified by IPT use are shown in Table 1.

**Figure 1.**
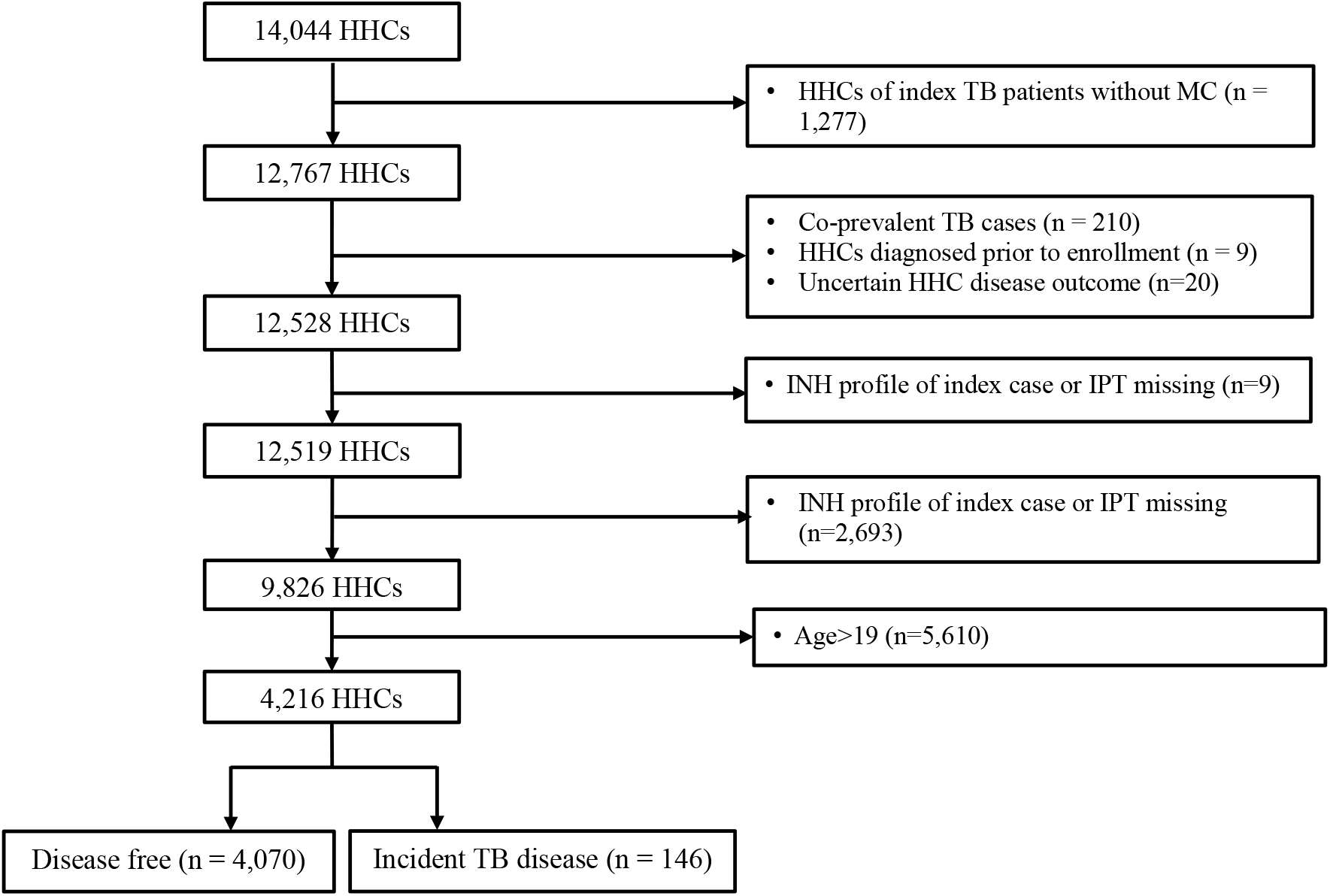
Flow diagram of household contacts of household contacts of index TB patients

**Table 1.** Baseline characteristics of household contacts with age ≤ 19, stratified by receiving isoniazid prevention therapy or not.

| Characteristic (total N with data) | Not received IPT |  | Received IPT |  |
| --- | --- | --- | --- | --- |
|  | N | % | N | % |
| Age years (N=4,216) |  |  |  |  |
| 0 to 5 | 664 | 31% | 855 | 41% |
| 6 to 10 | 439 | 21% | 532 | 25% |
| 11 to 15 | 489 | 23% | 451 | 22% |
| 16 to 19 | 528 | 25% | 258 | 12% |
| Male gender (N=4,216) |  |  |  |  |
| Female | 1,087 | 51% | 1,033 | 49% |
| Male | 1,033 | 49% | 1,063 | 51% |
| HIV seropositive (N=4,164) |  |  |  |  |
| No | 2,086 | 100% | 2,074 | 100% |
| Yes | 4 | 0% | 0 | 0% |
| Diabetes (N=4,202) |  |  |  |  |
| No | 2,111 | 100% | 2,087 | 100% |
| Yes | 2 | 0% | 2 | 0% |
| BCG scars (N=4,216) |  |  |  |  |
| 0 | 423 | 20% | 401 | 19% |
| 1 | 1,640 | 77% | 1,650 | 79% |
| $\geq 2$ | 57 | 3% | 45 | 2% |
| Smoking status (N=4,209) |  |  |  |  |
| Non-smoker | 2,068 | 98% | 2,086 | 100% |
| 1 cigarette per day | 25 | 1% | 5 | 0% |
| >1 cigarette per day | 22 | 1% | 3 | 0% |
| Alcohol use (N=4,195) |  |  |  |  |
| Non-drinker | 1,912 | 91% | 2,006 | 96% |
| 0 to <3 drinks per day | 149 | 7% | 73 | 4% |
| $\geq 3$ drinks per day | 44 | 2% | 11 | 1% |
| Nutritional status<sup>a</sup> (N=4,173) |  |  |  |  |
| Normal weight | 1,748 | 83% | 1,681 | 81% |
| Underweight | 44 | 2% | 59 | 3% |
| Overweight | 308 | 15% | 333 | 16% |
| Employed outside the home (N=4,214) |  |  |  |  |
| No | 1,893 | 89% | 1,981 | 95% |
| Yes | 226 | 11% | 114 | 5% |
| Use of public transportation (N=4,120) |  |  |  |  |
| Non-user | 736 | 35% | 795 | 39% |
| 1 to 3 days per week | 709 | 34% | 640 | 32% |
| 4 to 7 days per week | 652 | 31% | 588 | 29% |
| Socioeconomic status<sup>b</sup> (N=4,128) |  |  |  |  |
| Low | 821 | 40% | 801 | 39% |
| Middle | 931 | 45% | 887 | 43% |
| High | 325 | 16% | 363 | 18% |
| TB infected at baseline (N=4,068) |  |  |  |  |
| No | 1,417 | 70% | 1,494 | 73% |
| Yes | 613 | 30% | 544 | 27% |
| Index-case INH-profile (N=4,216) |  |  |  |  |
| Sensitive | 1,534 | 72% | 1,630 | 78% |
| Mono-resistant | 185 | 9% | 201 | 10% |
| MDR | 401 | 19% | 265 | 13% |

<sup>a</sup> Nutritional status was defined by the WHO body mass index z-score tables
<sup>b</sup> Socioeconomic status was defined using a principal component analysis based on housing quality, water supply, and sanitation.
Abbreviation: N: number; TB: tuberculosis; INH: isoniazid; IPT: isoniazid prevention therapy; MDR: multi-drug resistant

At 12–months follow–up, 146 HHCs developed TB disease. Of these, 48 (33%) had complete 24–loci MIRU–typing. Twenty–nine of the 48 (64%) had at least 23 loci that matched their index cases’ MIRU–typing.

### Univariate analyses and multivariate adjustment

HHCs under age 15 who received IPT were one third as likely to develop TB disease compared to those who did not in both the univariate and multivariate model adjusted for age, SES, and history of TB (HR=0·33, 95% CI = 0·22–0·48 and adjusted HR=0·34, 95% CI=0·23–0·50) (Table 2).

**Table 2.**
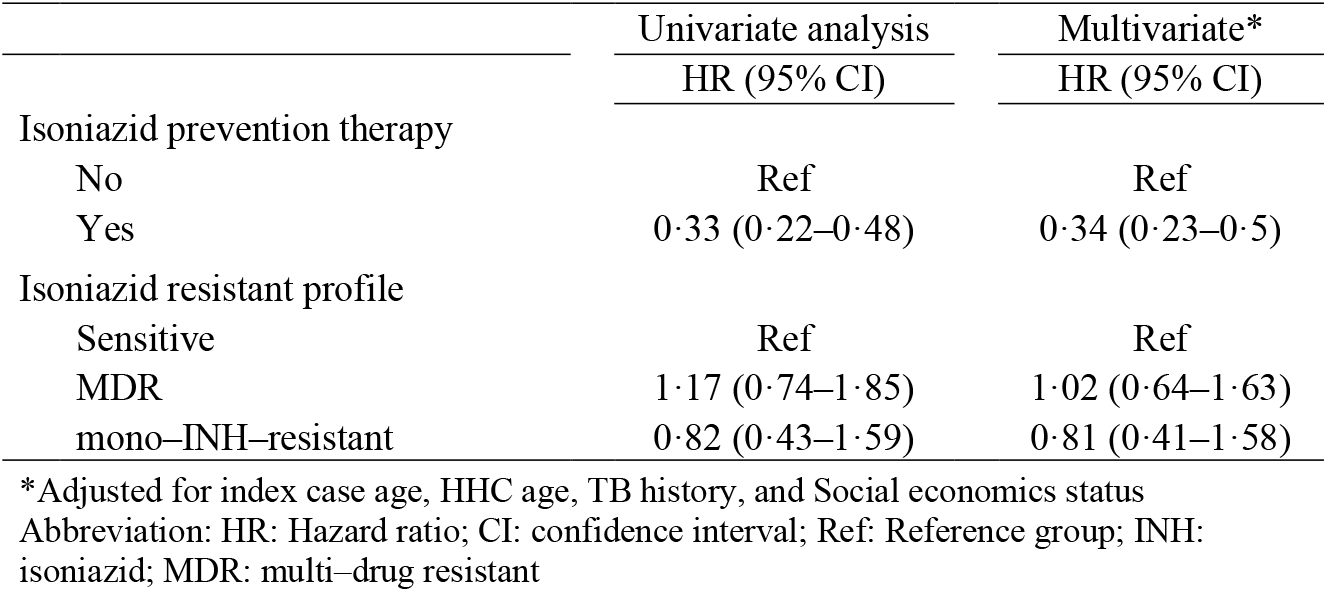
Univariate– and multivariate–adjusted effects of isoniazid prevention therapy, and the isoniazid resistant profile pattern of tuberculosis index cases on disease incidence of household contacts ≤ 19 years of age.

|  | Univariate analysis | Multivariate* |
| --- | --- | --- |
|  | HR (95% CI) | HR (95% CI) |
| Isoniazid prevention therapy |  |  |
| No | Ref | Ref |
| Yes | 0.33 (0.22–0.48) | 0.34 (0.23–0.5) |
| Isoniazid resistant profile |  |  |
| Sensitive | Ref | Ref |
| MDR | 1.17 (0.74–1.85) | 1.02 (0.64–1.63) |
| mono–INH–resistant | 0.82 (0.43–1.59) | 0.81 (0.41–1.58) |

\*Adjusted for index case age, HHC age, TB history, and Social economics status
Abbreviation: HR: Hazard ratio; CI: confidence interval; Ref: Reference group; INH: isoniazid; MDR: multi–drug resistant

### Adding IPT and INH–resistant profile of index case as interaction terms

INH effectiveness was higher in HHCs exposed to DS or MDR–TB than in those exposed to mono–INH–R strains (IPT vs. No–IPT adjusted HR=0·32, 95% CI=0·20–0·50 in INH–sensitive subgroup; 0·26, 95% CI=0·08–0·77 in MDR; 0·80, 95% CI=0·23–2·79 in mono–INH–resistant) (Table 3A). IPT effectiveness was increased further in the subgroup who received IPT for more than 3 months across all three resistance categories (adjusted HR=0·20, 95% CI=0·10–0·40 in INH–sensitive subgroup; 0·16, 95% CI=0·02–1·27 in MDR; 0·72, 95% CI=0·16–3·16 in mono– INH–resistant) (Table 3B) and reduced in those treated for less than three months (IPT vs. No– IPT adjusted HR 089, 95% CI=0·44–1·82 in INH–sensitive subgroup; 0·52, 95% CI=0·09–1·84 in MDR; 1 02, 95% CI=0·10–8·46 in mono–INH–resistant) (Table 3C). Among participants ≤ five years old, IPT effectiveness was almost 100% in those who received ≥ three months treatment (Table 4A–4C). Among HHCs for whom index patient minimal inhibitory concentrations (MICs) were available (N=1,276), IPT efficacy remained high among HHCs exposed to INH–sensitive and INH–moderate phenotypes (MIC ≤5μg/ml) (IPT vs. No–IPT HR=0·37, 95% CI=0·10–1·37). Among 92 HHCs who received IPT after being exposed to an index patient with an MIC >5 μg/ml, none developed (0/92) active TB, while 4% (14/368) of those who did not receive IPT developed disease.

**Table 3.** The effect of isoniazid prevention therapy on disease incidence of children ≤19 years of age, stratified by INH profiles of index cases; adjusted for index case age, HHC age, TB history and social economics status.

|  | Isoniazid-sensitive<br>N=3,099; event=106 | MDR<br>N=664; event=27 | Mono-isoniazid resistant<br>N=365; event=11 |
| --- | --- | --- | --- |
| INH prevention therapy | HR (95% CI) | HR (95% CI) | HR (95% CI) |
| No | Ref | Ref | Ref |
| Yes | 0.32 (0.2–0.5) | 0.26 (0.08–0.77) | 0.8 (0.23–2.79) |
| Likelihood ratio test for interaction term: 0.015 |  |  |  |

B. Household contacts who received isoniazid prevention therapy $\geq 3$ months
|  | Isoniazid-sensitive<br>N=2,429; event=88 | MDR<br>N=505; event=24 | Mono-isoniazid resistant<br>N=299; event=9 |
| --- | --- | --- | --- |
| INH prevention therapy | HR (95% CI) | HR (95% CI) | HR (95% CI) |
| No | Ref | Ref | Ref |
| Yes | 0.2 (0.1–0.4) | 0.16 (0.02–1.27) | 0.72 (0.16–3.16) |
| Likelihood ratio test for interaction term: <0.001 |  |  |  |

C. Household contacts who received isoniazid prevention therapy < three months
|  | Isoniazid-sensitive<br>N=1,727; event=88 | MDR<br>N=465; event=25 | Mono-isoniazid resistant<br>N=210; event=7 |
| --- | --- | --- | --- |
| INH prevention therapy | HR (95% CI) | HR (95% CI) | HR (95% CI) |
| No | Ref | Ref | Ref |
| Yes | 0.89 (0.44–1.82) | 0.52 (0.11–2.38) | 1.02 (0.11–9.4) |
| Likelihood ratio test for interaction term: 0.367 |  |  |  |

### Second independent dataset

The second study included 1,121 HHCs ≤ 19 years age who had available IPT data. Here, again, IPT use was associated with reduced rates of incident TB in both univariate and analyses that adjusted for age, SES, and TB history (HR=0·1; 95% CI=0·03–0·30 and adjusted HR=0·11; 95% CI=0·02–0·49). IPT not only protected HHCs of drug–sensitive index cases (adjusted HR=0·13 95% CI=0·03–0·57), but none of 76 HHCs of MDR–TB index cases who received IPT developed TB compared to 8/273 (3%) without IPT.

## Discussion

Here, we found that IPT is associated with reduced incidence of TB disease among HHCs of TB patients even when the index patients were infected with MDR and mono–INH–resistant Mtb strains. INH effectiveness was higher among HHCs of MDR–TB compared to mono–INH resistant strains. As expected, TB risk was higher in those who received less than three months therapy, especially among children under five years of age. No child who received at least three months of IPT developed TB disease. We also showed that the effectiveness of IPT is unrelated to the MIC of the index patient’s TB strain; no HHC who was exposed to an index patient with a >5 μg/ml MIC developed disease.

Few data exist on the effectiveness of INH in preventing TB progression among people exposed to MDR–TB. In Brazil, Kritski et al., investigators followed 190 TST–positive contacts of MDR–TB patients and found that disease developed in two of 45 (4%) contacts who received IPT and in 13 of 145 (9%) contacts who did not.^12^ In Israel, Attamna et al. followed contacts of MDR–TB cases for up to six years and reported no cases among 71 contacts receiving IPT, and suggested that IPT might have been effective in preventing the progression of MDR–LBTI.^13^ In South Africa, Schaaf et al. reported that children who did not receive preventive therapy were four times more likely to develop TB disease than those who received less than six months of individualized preventive therapy that contained a high dose of INH (15–20 mg/kg/d).^14^ Although this suggests that INH could have been effective against MDR–LTBI, its effectiveness could not be measured as the regimens contained other drugs tailored to the drug susceptibility profile of the index strain. A study conducted in Australia compared people exposed to MDR–TB who received IPT to those who received other preventive therapy regimens or no treatment.^15^ Of these, two contacts developed TB disease within 54 months, but the study did not specify what regimens these two incident patients received. Other studies which reported on regimens that included INH among contacts of MDR–TB patients lacked control arms.^16–18^

We considered several possible explanations for the observed effectiveness of IPT among contacts of INH–resistant TB patients. First, HHCs might have been infected in the community by patients with drug–sensitive TB rather than the drug–resistant TB patient living in their households. However, approximately two–thirds of the patients with secondary disease harbored strains that matched their index case, suggesting that no more than a third of the secondary cases acquired TB in the community. If IPT had no effect in preventing MDR TB, we would expect that this potential misclassification would lead to an observed protective effect of no less than 0·32 (effect size of contacts exposed to drug sensitive strains, Table 3A), rather than the 0·26 effect we found. Furthermore, the observed effect was more extreme in under–5 year olds, whom we considered much less likely than older contacts to have been infected by someone other than the index case. In the independent dataset, Grandjean et al. noted that 86% of MDR–TB secondary cases were exposed to MDR–TB index cases, again suggesting that most of the incident MDR–TB among HHCs in their study were infected at home rather than in the community.^10^

Secondly, we considered the possibility that HHCs who chose to take IPT came from higher SES groups and thus were less likely to develop TB disease regardless of the resistance profile of the index case. Although we attempted to adjust for SES, it is possible the principal component score we used did not completely capture its effect. However, in this case, we would not expect the INH effect to vary by duration of therapy as it did in our study (Table 3 and 4). The reduced efficacy of IPT among people who received less than one month of treatment is within the range reported in a highly referenced randomized trial, again suggesting that confounding introduced by SES could not explain our findings.^10^

**Table 4.**
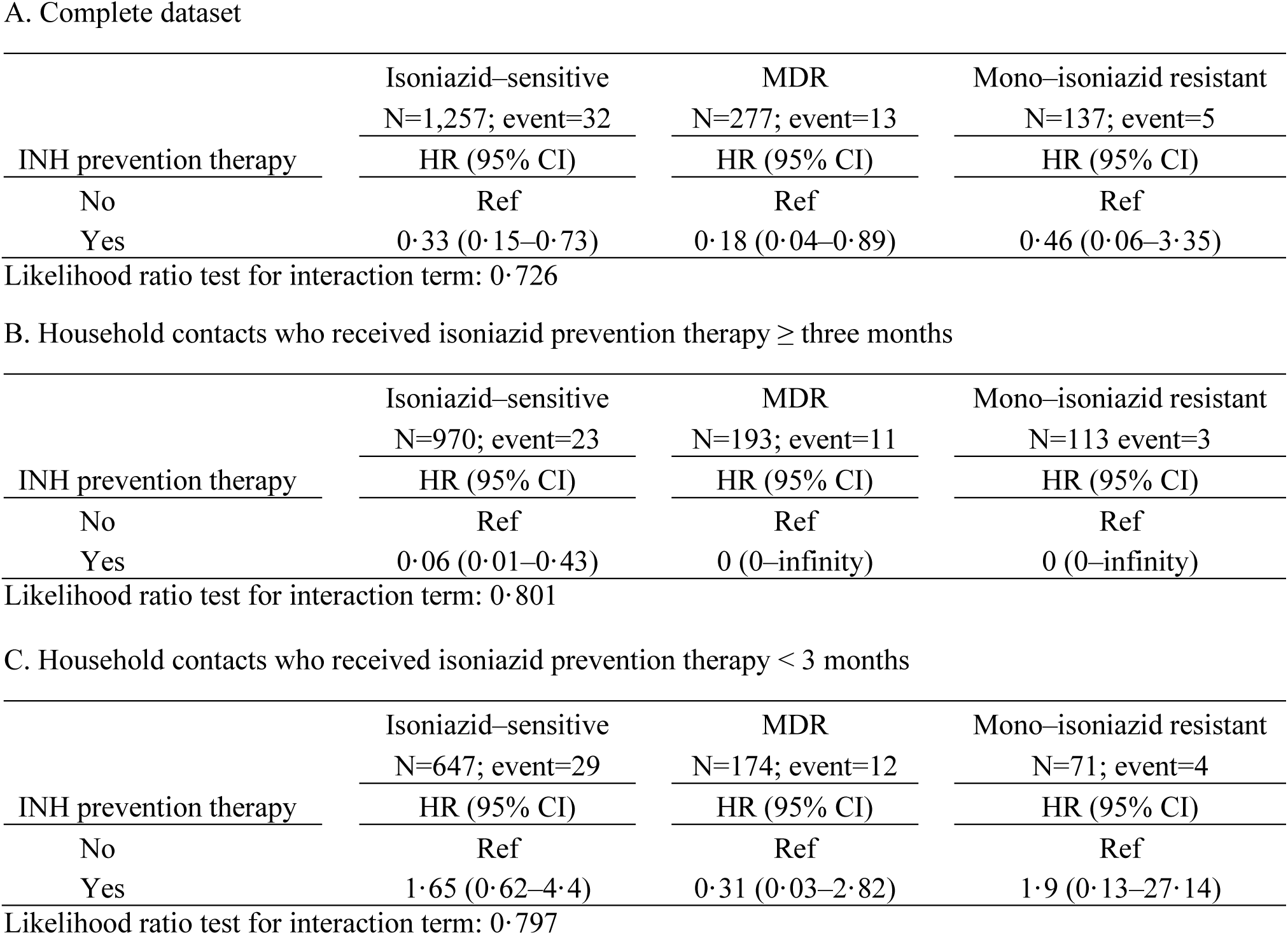
The effect of isoniazid prevention therapy on disease incidence of children ≤ five years of age, stratified by INH profiles of index cases; adjusted for index case age, HHC age, TB history and social economics status.

Finally, we considered the possibility that INH is effective against LTBI even when the relevant strains are found to be resistant to INH in media–based growth assays. This raises the possibility that the mechanism by which INH reduces TB risk among those with LTBI may differ from its mechanism in TB disease. In the latter case, INH is known to be a pro-drug which is converted to its active metabolite, an INH–NAD adduct, by an MTB catalase peroxidase encoded by the KatG gene.^19^ The INH–NAD adduct then binds to InhA (enoyl–acyl carrier protein reductase) and inhibits the synthesis of essential mycolic acids in MTB cell walls. The most common causes of INH resistance among clinical strains are mutations in KatG that reduce the activity of the catalase-peroxidase and thereby block the conversion of INH to its active form. Several studies have raised the possibility that this conversion may occur independently through other routes. Youatt et al. showed that the presence of copper increased the INH sensitivity of an INH-resistant strain, suggesting the interaction of INH and copper ions may facilitate the conversion of INH to its active form.^19,20^ In a second study, Mahapatra et al. identified metabolites of oxidized INH-NAD adducts in the urine of people who were not infected with MTB, thereby demonstrating that INH can be activated by host enzymes.^21^ Other studies have suggested that INH may employ nonspecific antibacterial mechanisms against MTB in addition to its impact on mycolic acid synthesis. INH is a strong ligand for iron, copper and zinc and might be involved in metal ion uptake by MTB, which could disrupt metal homeostasis and inhibit MTB growth.^22–24^

These hypotheses raise the question of why INH fails to cure INH–resistant TB patients. One possible explanation is that these mechanisms clear MTB in the early stage of infection when the bacterial load is low, but are less effective when the bacterial load is much higher. Another explanation is that INH may kill latent TB through other unaddressed mechanisms, as several studies have hypothesized that latent TB is in a cell-wall deficient form and so INH cannot have a bactericidal effect through the inhibition of cell wall synthesis.^25,26^

Our study also showed that the protective effect of INH differs in contacts exposed to MDR-TB strains compared to mono–INH–resistant strains. While this could be due to random variation related to the small sample size of HHCs exposed to mono–INH–resistant TB, another possibility is suggested by the finding that INH mutation profiles differ between MDR and mono–INH–resistant strains. Alland et al. reported that mono–INH–resistant strains were more likely than MDR strains to harbor InhA promoter mutations and less likely to have KatG mutations.^27^ Since InhA is the downstream target of the INH–NAD adduct, mono–INH–resistant strains may remain resistant to INH regardless of whether INH conversion took place through an MTB–dependent or independent pathway.

IPT has been used for decades in tuberculosis control efforts and despite some concerns about hepatoxicity, it has been shown to have a good safety profile especially in children. Health workers worldwide have extensive experience using this drug and handling its adverse effects. Establishing its efficacy against latent MDR TB would therefore be of great value and could set a bar against which alternative treatment could be measured. For example, the ongoing PHOENIx trial, designed to establish the efficacy of delamanid against MDR-LTBI uses INH as the control arm.^28^ If the investigators of PHOENIx trial consider INH is ineffective against MDR-LTBI and is serving only as a placebo, the efficacy of delamanid against MDR-LTBI could be underestimated.

Our study has some limitations. The contacts of MDR–TB cases received INH for a shorter period of time than contacts of pan–sensitive or mono–INH–resistant cases, presumably because clinicians halted IPT once the index patients’ MDR–TB status were confirmed. Given the dose effect we observed, we would expect to see an even more extreme effect of IPT had contacts of MDR–TB cases received the same duration of IPT as those exposed to drug–sensitive strains. Also, we were unable to assess the effect of IPT on adult contacts of MDR–TB cases given that IPT is not indicated for adult contacts without co–morbidities in Peru. Finally, almost all HHCs in our cohort were HIV–negative, so we were not able to evaluate the synergistic effect between IPT and highly active antiretroviral therapy in HIV–positive HHCs exposed to MDR–TB.

In conclusion, we found that IPT protected against TB among contacts of INH–resistant TB patients. Given the safety profile of INH and its wide use across the globe, INH may have a role in the management of MDR–LTBI.

## Author Contributions

MBM and MCB led the study design. LL oversaw data collection and management with JTG, RC, ZZ, CC, and RY. RC managed laboratory efforts. MBM supervised data analysis and interpretation in conjunction with C-CH and MFF. C-CH and MBM wrote the first draft of the manuscript, and all authors contributed to manuscript revision.

## Declaration of Interests

Authors declare no competing interests.

## Acknowledgments

We thank the patients and their families who gave their time and energy to contribute to this study, the National Strategy for TB Control at the Peruvian Ministry of Health, and the healthcare personnel at the 106 participating health centers in Lima, Peru. This work was supported by the National Institutes of Health and the National Institute of Allergy and Infectious Diseases CETR [U19AI109755] and TBRU [U19AI111224].

