## supplement 1 for "The impact of isoniazid preventive therapy on tuberculosis among household contacts of isoniazid–resistant patients"

### Recruitment

This study was conducted in Lima in 106 district health centers that provide care to a population of approximately three million residents. Patients were referred to study staff if they were over 15 years of age and had been diagnosed with pulmonary TB (PTB) disease by a health center clinician on the basis of sputum smear microscopy or chest radiography. We collected an additional sputum sample from consenting participants which we sent for repeat sputum smear microscopy, mycobacterial culture, and drug sensitivity testing. We requested permission to visit each patient's household and recruit his or her household contacts (HHCs) into a prospective cohort study. Study workers aimed to enroll all household members within one week of the diagnosis of the index case.

### Baseline assessment of index patients

We collected the following data from index patients at the time of enrollment: age, gender, occupation, symptoms of TB, duration of symptoms, history of TB disease, alcohol, intravenous drug and tobacco history, and comorbidities including HIV and diabetes mellitus. Patients who did not know their HIV status had blood drawn for HIV and CD4 count. Signs associated with TB disease, height, and weight were recorded. Index patients also underwent HIV testing and were evaluated with a chest radiograph. The time to treatment was measured as the number of days the patient reported coughing prior to diagnosis.

### Bacteriological cultures and drug susceptibility testing

Sputum samples were tested for the presence of acid–fast bacilli by Ziehl–Neelsen staining and cultured by inoculation in two tubes containing Lowenstein–Jensen or Ogawa medium. Indirect susceptibility testing to INH (INH), Rifampicin (RIF), Ethambutol (EMB) and Streptomycin (STR) was conducted by the Löwenstein–Jensen Proportion Method, using the following drug concentrations: INH (0·2 and 1·0 µg/ml), RIF (40·0 µg/ml), EMB (2·0 µg/ml), and STR (4·0 µg/ml). Susceptibility to Pyrazinamide (PZA) (100 ug/ml) was tested using the Wayne method. DNA from each mycobacterial culture was extracted and genotyped by 24–loci mycobacterial interspersed repetitive units–variable–number tandem repeats (MIRU–VNTR) using standard methods.[1]

### Follow–up of index patients

Index patients received directly observed therapy at their district health clinics, as specified in the Peruvian National Tuberculosis Control Program (NTP) guidelines for drug–sensitive and drug–resistant TB. Patients with drug–sensitive TB received a standard 6–month course with a 2–month "intensification phase" of INH, RIF, PZA, and EMB followed by a 4–month "consolidation phase" of INH and RIF alone. Patients with MDR–TB received treatment according to NTP guidelines. Since results for routine drug resistance testing were often not available for two to three months after initial diagnosis, patients who were not previously suspected of having MDR–TB were started on a first–line drug regimen until MDR–TB was confirmed.

### INH preventive therapy for HHCs

The 2006 Peruvian National TB Program recommended that HHCs 19 years old or younger and those who had a specified comorbidity should receive six months of IPT while those with HIV should receive 12 months.[2] Children aged 19 and under were offered IPT at the time index patients were diagnosed, regardless of TST status. Health care providers often chose to discontinue IPT in HHCs if the index patient was subsequently diagnosed with MDR–TB but some MDR–exposed HHCs received a full course of IPT. We used medical records from participating hospitals and health clinics to determine the duration of IPT.

### Enrollment of household contacts

At the time of the enrollment of HHCs, study workers collected the following data: whether IPT had been initiated, age, gender, relationship to index patient, housing information including number of rooms, building material, type of flooring, income, education, history of incarceration, occupation, alcohol, cigarette and illicit drug intake, general health history including previous history of TB, BCG vaccination, co–morbidities, BMI and medications taken. Participants were assessed for symptoms associated with TB disease including cough, night sweats, weight loss, and fever. Those with symptoms were referred to their local health clinic for chest radiography and clinical evaluation for active TB disease. Household members with no known history of active TB disease or previously documented infection received a tuberculin skin test (TST), and those with unknown HIV status were tested for HIV.

**Follow–up of household contacts**

Participants were revisited in their household at two, six, and 12 months and were asked whether they had been diagnosed with TB or if they had had symptoms of active disease. Those who reported symptoms were referred to their local health center for further clinical evaluation including a chest radiograph and sputum smear. Participants who tested negative at the initial study visit and who had not developed active TB disease at the time of the follow–up visit underwent repeat TST and clinical evaluation at 6sixand 12 months.  We used medical records from participating hospitals and health clinics to determine the duration of IPT.

**Data categorization**

We considered HHCs to have received IPT in response to the exposure to the index patient if INH was initiated within three months of that patient's diagnosis. We categorized participants according to their alcohol intake as nondrinkers if they reported having consumed no alcoholic drinks per day, light drinkers if they reported drinking <40 grams or <3 alcoholic drinks per day, and heavy drinkers if they reported drinking 40 grams or more of alcohol or three or more drinks per day. A large proportion of smokers reported smoking only a single cigarette per day. We classified people as nonsmokers if they reported no cigarette smoking, as light smokers if they reported smoking one cigarette per day, and as heavy smokers if they reported smoking more than one cigarette per day. We defined nutritional status for children based on the WHO body mass index (BMI) z–score tables.[3] We assigned people with BMI z–scores of less than two as underweight and those greater than two as overweight.

We created a continuous variable to capture household socioeconomic status (SES) by including variables on housing quality, water supply, and sanitation in a principal component analysis (PCA). PCA is a data reduction statistical technique that extracts a set of uncorrelated 'principal components' from a set of correlated variables, where each principal component is a weighted linear combination of the original variables. The continuous SES score was categorized into tertiles corresponding to relative "low," "middle," and "upper" SES.

**Outcome definition**

We identified incident TB among HHCs during scheduled household visits and from a systematic review of TB registries at the participating health clinics. We considered HHCs to have co–prevalent TB if they were diagnosed within two weeks of the diagnosis of the index case. If HHCs were diagnosed between two weeks and 15 months after diagnosis of the index case, we considered them "secondary" cases. Diagnosis of adult secondary TB followed the same criteria as outlined above for index cases. We defined secondary TB disease among contacts younger than 18 years of age according to the consensus guidelines for classifying TB disease in children.[4]

**Analyses**

We included in our analysis only HHCs under 19 because older contacts were only offered IPT if they had comorbidities that substantially increased their risk of TB disease.  We used a Cox frailty proportional hazards model to evaluate risk factors for incident TB disease, accounting for clustering within households.[5] We first performed a univariate analysis to examine the effect of IPT on TB incidence, followed by a multivariate model in which we adjusted for the age of the index case age and the age, SES and TB history of the HHC. To evaluate whether the effect of IPT on TB incidence varied by resistance profile of the index case, we added a variable representing INH resistance in the index case and an interaction term for INH–resistance and IPT. Because the spectrum of INH resistance–causing mutations that lead to INH mono–resistance may differ from those that lead to MDR–TB, we classified strains as sensitive, mono–INH–resistant, or MDR–TB (resistant to both INH and RIF). Previous studies have shown that the efficacy of IPT treatment is reduced if the treatment is ended within three months.[6] We therefore repeated these analyses stratifying by a dichotomous variable that captured treatment for more or less than three months. We also considered the possibility that HHCs ≤ 5 years of age would be more likely to acquire TB at home than in the community compared to older contacts and we thus conducted sensitivity analyses restricted to this subgroup.

To determine whether the effect of IPT on disease in the HHCs was a function of the mean inhibitory concentrations (MICs) of the infecting organism, we repeated these analyses for the subset of HHCs exposed to index cases for whom quantitative INH–resistance was available.

**Verifying our finding with an independent dataset**

We conducted a similar analysis using publically available data from an independent dataset collected from a prospective cohort study in South Lima and Callao, Peru between 2010 and 2013, posted by Grandjean et al.[7] This study enrolled 1,055 HHCs of 213 MDR–TB index cases and 2,362 HHCs of 487 drug–susceptible index cases and measured incident TB over 2–years of follow–up. Drug susceptibility testing for INH and RIF was performed for all index cases' samples using microscopic observation drug susceptibility assays in regional laboratories and results were confirmed in the national reference laboratory using proportions methods.[8] The investigators note that IPT was discontinued in this group after MDR–TB index cases were confirmed but data on the duration of IPT were not available.

We used a Cox frailty proportional hazards model to evaluate the association between IPT and incident TB infection in individuals aged 19 and under, accounting for clustering within each matched set. We first performed univariate analysis, followed by a multivariate model adjusted for HHCs' age, SES, and previous TB history. We then added a dichotomous variable for the drug resistance status (MDR or sensitive) in the index case, as well as interaction terms for the resistance profile and IPT to evaluate whether the effect of IPT on TB incidence varied by the resistance profile of index cases.
